# A Novel Tool for the Removal of Muscle Artefacts from EEG: Improving Data Quality in the Gamma Frequency Range

**DOI:** 10.1101/2020.11.23.393702

**Authors:** Alina Pauline Liebisch, Thomas Eggert, Alina Shindy, Elia Valentini, Stephanie Irving, Anne Stankewitz, Enrico Schulz

## Abstract

**Background:** The past two decades have seen a particular focus towards high-frequency neural activity in the gamma band (>30Hz). However, gamma band activity shares frequency range with unwanted artefacts from muscular activity.

**New Method:** We developed a novel approach to remove muscle artefacts from neurophysiological data. We re-analysed existing EEG data that were decomposed by a blind source separation method (independent component analysis, ICA), which helped to better spatially and temporally separate single muscle spikes. We then applied an adapting algorithm that detects these singled-out muscle spikes.

**Results:** We obtained data almost free from muscle artefacts; we needed to remove significantly fewer artefact components from the ICA and we included more trials for the statistical analysis compared to standard ICA artefact removal. All pain-related cortical effects in the gamma band have been preserved, which underlines the high efficacy and precision of this algorithm.

**Conclusions:** Our results show a significant improvement of data quality by preserving task-relevant gamma oscillations of cortical origin. We were able to precisely detect, gauge, and carve out single muscle spikes from the time course of neurophysiological measures. We advocate the application of the tool for studies investigating gamma activity that contain a rather low number of trials, as well as for data that are highly contaminated with muscle artefacts. This validation of our tool allows for the application on event-free continuous EEG, for which the artefact removal is more challenging.

## Introduction

Electrophysiological research spanning more than 20 years yielded substantial evidence that high-frequency neural activity in the gamma band is involved in neuronal signal transmission (Cardin et al., 2009; Knoblich et al., 2010; Michalareas et al., 2016), as well as for sensory, perceptual, cognitive and motor processing in the brain (Bertrand and Tallon-Baudry, 2000; Debener et al., 2003; Tallon-Baudry et al., 2005).

However, the interpretation of this evidence is challenged by major sources of signal of extra-neural origin, e.g. voluntary and autonomic muscle activity (Muthukumaraswamy, 2013; Whitham et al., 2007), eye movements (Jung et al., 2000), saccades (Carl et al., 2012; Hassler et al., 2011; Keren et al., 2010), as well as environmental electrical noises (de Cheveigné, 2020). While power line artefacts and eye movements can be successfully detected, the contamination of electroencephalography (EEG) with muscle activity has not yet been sufficiently addressed. The problem is that the spectral bandwidth of muscle activity coincides with neural activity in the gamma band (Hipp and Siegel, 2013; Muthukumaraswamy, 2013).

Distinct muscle artefacts can be removed by using spatial filtering such as independent component analysis (ICA; Delorme et al., 2007; Ma et al., 2012; Shackman et al., 2009) and the inspection of the single trial time-frequency decomposed ICA data (Michail et al., 2016; Schulz et al., 2011). However, this procedure entirely relies on the researcher’s expertise and casts some degree of arbitrariness in adjudicating whether an independent component (IC) or single trial contains artefacts or not. This is particularly true for ICs that exhibit occasional muscle spikes and yet may also contain cortical activity. However, the decision as to whether ICs can be removed is relatively straightforward in event-related designs, for which the induced response of each IC can be evaluated (Michail et al., 2016). In contrast, such decisions are more difficult for ongoing neuronal processes or investigations of resting-states. In the absence of any event, these present the researcher with the issue that for some ICs, minimal muscle-generated activity can temporarily co-occur with cortical activity.

Particular attention is needed for contaminating muscle activity in studies that require eye movements to execute the experiment (Yuval-Greenberg et al., 2008), as well as for studies using experimentally-applied pain. Phasic and tonic noxious stimuli can cause stimulus-related saccades and muscle twitches, and continuous muscle activity decreases the signal-to-noise ratio (SNR) in the gamma range. Unfortunately only a few studies on pain that investigate laser-evoked gamma activity conduct an inspection of single trial time-frequency data (Liberati et al., 2018; Michail et al., 2016; Schulz et al., 2012). This is of utmost importance as a few stimulus-related muscle twitches, whose amplitude is substantially larger than cortical gamma, can contribute to the induced potential of the subject and can, in the case of low sample sizes, contribute to the study result. Indeed, some studies on pain have revealed gamma effects with an unusually broad frequency range (Zhang et al., 2012) or scalp distribution (Chien et al., 2014), which suggests contamination of the results with muscle activity.

Therefore, we developed a novel tool that is able to detect and to remove isolated single muscle spikes whilst leaving the remaining time course of cortical activity intact. Specifically, we provide evidence that this tool is capable of mending slightly contaminated ICs and trials.

## Materials and methods

### Subjects

Raw data were taken from our previous study (Michail et al., 2016). EEG was recorded from nineteen healthy male human subjects with a mean age of 24 years (21–31 years). All subjects gave written informed consent. The study was approved by the local ethics committee and conducted in conformity with the Declaration of Helsinki.

### Paradigm

Seventy-five painful cutaneous laser stimuli were delivered to the dorsum of the right hand. The laser device used was a Tm:YAG laser (Starmedtec GmbH, Starnberg, Germany) with a wavelength of 1960 nm, a pulse duration of 1 ms, and a spot diameter of 5 mm. The physical energy of the painful stimulation was kept constant at 600mJ. To prevent skin damage, the stimulation site was changed slightly after each stimulus.

The interstimulus interval (ISI) was randomly varied between 8 – 12 s. To prevent excessive eye movement-related artefacts, the subjects perceived the stimuli with closed eyes. Three seconds after each stimulus, the subjects were prompted by an auditory cue to verbally rate the perceived intensity of the pain stimulus on a 0–10 numerical rating scale, where 0 indicated no pain and 10 the most intense pain imaginable.

### EEG Recordings

EEG data were recorded using an MRI-compatible electrode cap (FMS, Munich, Germany). The electrode montage included 64 electrodes consisting of all 10–20 system electrodes and the additional electrodes Fpz, FCz, CPz, POz, Oz, Iz, AF3/4, F5/6, FC1/2/3/4/5/6, FT7/8/9/10, C1/2/5/6, CP1/2/3/4/5, TP7/8/9/10, P5/6, PO1/2/9/10, plus two electrodes below the outer canthus of each eye. The EEG was referenced to the FCz electrode, grounded at AFz, sampled at 1 kHz with 0.1 μV resolution. Impedance was kept below 20 kΩ.

### EEG preprocessing

Raw EEG data were preprocessed in Brain Vision Analyzer Software (Brain Products, Munich, Germany). After application of a 0.5 Hz high-pass filter, data were decomposed into 64 components using ICA. Components related to horizontal and vertical eye movements, as well as major artefacts, were removed and data were back-transformed into EEG signals. In order to compare the effect of the muscle spike tool, we processed the data in two parallel branches: (1) without, and (2) with muscle spikes removal:

(1) *Processing branch without muscle spike removal.* A further ICA was computed, where the number of components was aimed to be kept identical for all data and therefore limited to 55. ICA time courses were epoched around stimuli between −1.55 and +2 s and exported to Matlab (The Mathworks, USA), time-frequency decomposed, and z-transformed. A visual inspection on the single trial images of time-frequency representations (TFR) was performed. Artefact components were removed and the data were then back-projected to EEG channels.
(2) *Processing branch with muscle spike removal.* A further ICA specifically for the removal of muscle spikes on the time course of the ICs was computed. The number of components was aimed to be kept identical for all data and therefore limited to 55. The ICA decomposed data were epoched around stimuli between −1.55 and +2 s and exported to Matlab. In component space, correction of muscle artefacts was performed with a custom programme written in Matlab to identify and eliminate gamma artefacts (for details of the spike removal algorithm see below). For a quality check of the procedure, the cleaned data were time-frequency decomposed and z-transformed. A visual inspection on the single trial TFR images was performed. In case of satisfying cleaning, IC time courses were back-projected to EEG. A final ICA was performed in order to remove remaining artefact components.
(1&2) *Further analysis for both types of preprocessing branches.* Epoched artefact-free data of each EEG electrode were downsampled to 512 Hz, re-referenced to the common average reference, and time-frequency decomposed (for details see Michail et al., 2016). The single trials were baseline corrected by subtracting the pre-stimulus period (−1000 to −100 ms). Trials that still contained artefacts were also removed (for details and examples of artefacts in single trial representations (see Michail et al., 2016).

### Artefact correction using despiking

A very important aspect of our software is the automatisation of the artefact identification on ICA data. A previous study (Nottage et al., 2013) identified saccade-related muscle potential by localising peaks in the second derivative of the EEG signal that occurred simultaneously with acceleration peaks in the EOG channel. This method is suitable for eliminating saccade-related muscle artefacts, but can not remove artefacts caused by activity of scalp muscles. A major problem is that the artefact-related peaks in the EEG signal or its derivatives do not show statistics which are constant over time. Consequently, it is rather difficult to define absolute thresholds for their detection. Therefore, we developed an algorithm for analysing the distribution of the peak-to-peak amplitudes of fast slopes in the EEG signal. The algorithm works on a single time sequence and identifies muscle artefacts as large outliers in this distribution. It implements a cluster analysis of the local peak-to-peak amplitudes. The algorithm is applied in a sliding window and is therefore able to adapt to temporal variations of the distribution. The procedure of the cleaning process is as follows:

1. The signal for each ICA component was filtered with a median filter with length 0.1 s (median_filter_duration) and the output *(y_hp_*) of this lowpass was subtracted from the original signal. This “median highpass” was used to eliminate low frequencies.
2. Local peaks were detected in the filtered signal by defining a threshold which split the entire signal into sections where the signal was above or below the threshold. For each of the over-threshold-sections, a peak was defined by the time when the signal reached its maximum. The appropriate threshold was found from 50 potential values equidistantly distributed between the minimum and the maximum of the signal within the entire window. For each of these potential threshold values, it was counted how often the signal crossed, then the threshold with the largest number of crossings was selected (Fig. 1A, dashed light blue lines). This procedure was repeated for all window positions in increments of ¾ of the window duration. Afterwards, all local minima of the highpass-filtered trace were also localised by the same procedure applied on the flipped (i.e. multiplied by −1) trace.
3. From all local peak pairs closer to each other than 0.015 s (min_tpeak_diff), the peak with the smaller absolute values was eliminated until all remaining local peaks had a distance of more than 0.015 s (min_tpeak_diff) from its nearest neighbour.
4. Each of these local peaks was associated with a so-called *event neighbourhood* and an *event amplitude.* The event neighbourhood (see Fig. 1C) was defined by a window with the duration 0.2 s (cutoff_window_width) centred around each peak. If two or more neighbouring peaks fell in between the current peak and the upper or lower border, the respective border was adjusted to the centre between the two next-neighbour peaks. Also, the upper or lower border of the local neighbourhood was moved even closer to the peak if the highpass-filtered trace showed three or more zero-crossings between the peak and the respective border. In that case, the border was adjusted to the third zero-crossing. After determining the event neighbourhood, the event amplitude was defined by the difference between the maximum and the minimum of the highpass-filtered trace in the event neighbourhood of each peak.
5. All peaks with an event neighbourhood longer than a predefined limit (max. 0.025 s to 0.05 s; max_event_width) were excluded from further artefact analysis.
6. The remaining event amplitudes were submitted to a cluster analysis (Fig. 1B). The complete agglomerative hierarchical cluster tree was created (Matlab function *linkage)* and subsequently split into a fixed number (Nc) of clusters (Matlab function *cluster* with arguments ‘maxclust’, Nc). The number of clusters was determined as the one for which the distance of the median between the largest and the second-largest cluster was maximal (for 2≤ *Nc* ≤6). All peaks belonging to that largest cluster (as defined in step 4) were considered to be caused by a muscle artefact.
7. Steps 2 to 6 were executed within the same moving window. The window duration was set between 1 s and 40 s (analysis_window_width) and the window position increment was ¾ of the window duration.

**Figure 1:**
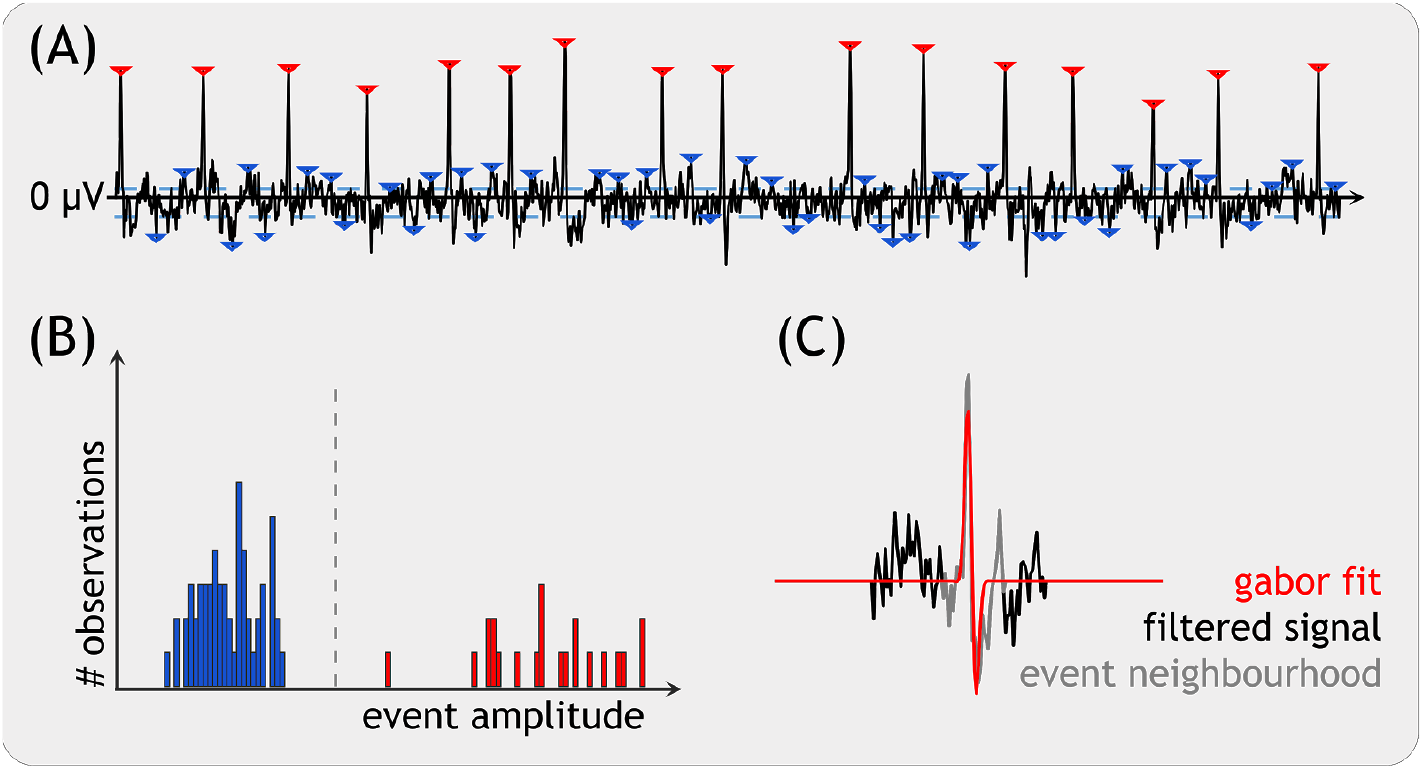
The figure illustrates the peak detection and spike removal for an exemplary analysis window. (A) Local maxima are defined as peaks if they exceed the threshold (dashed light blue line). The threshold line has the maximal possible number of line crossings from the filtered IC time course. The rectified super-threshold peaks are then submitted to a cluster analysis. (B) The peaks from the highest cluster (in red) are considered as artefact. The cluster threshold is shown as a vertical dashed line. (C) In order to estimate the size of the muscle spike a Gabor function was fitted into the signal.

The elimination of the artefacts identified with the above algorithm was then performed with a method very similar to the one previously utilised (Nottage et al., 2013). The highpass-filtered EEG signal (*y_hp_*, see step 1) was approximated by a Gabor function, within the neighbourhood of each artefact-peak (as defined in step 4). The Gabor function was defined as follows:

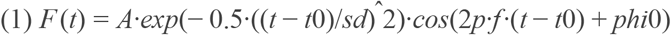

The five parameters [*A*, *t_0_, f*phi_*0*_, *sd*] were fitted by minimising (Matlab function fmincon) the squared difference between the Gabor function and *y_hp_* in two steps. In the first step, *A, f,* and phi_*0*_ were fitted while fixing *sd* to infinity and *t_0_* to the centre of mass of the absolute value of *y_hp_*. In the second step, all five parameters were fitted while using the results of the first fit as the initial value (Fig. 1C). Elimination was done by subtracting the fitted Gabor function from the original (non-filtered) EEG component trace.

### Statistical Analysis

Based on our previous findings (Michail et al., 2016; Schulz et al., 2012), we defined the gamma windows between 76-86 Hz and 0.15-0.35 s. Gamma responses were computed by subtracting a baseline (−1000 to −100 ms) for each trial (Hu et al., 2014). To compare both artefact cleaning approaches, paired t-tests were computed on the averaged trials and separately for each electrode. Additionally, we computed linear mixed effects models (LMEs) to explore the strength of the relationship between single trial neuronal responses and the respective pain ratings (Witkovský, 2012). The expected value of the response variable “rating” is modelled by a linear (regression) function that depends on the explanatory variable gamma activity. The LMEs were computed separately for each electrode:

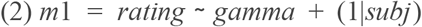

A further set of LMEs were computed to directly compare the relationships between both analysis strategies. This comparison aimed at elucidating whether the additional data cleaning exhibits a stronger relationship between neuronal responses and perception:

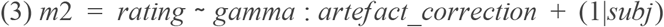

The resulting t-values of the fitted models are related to the estimated coefficient for the fixed effects (for the models within one modality) or to the interaction coefficients “gamma:artefact:correction” (for the models comparing both modalities), respectively. False discovery rate correction (FDR, Genovese et al., 2002) was used to correct for multiple comparisons.

## Results

### Artefact correction performance

The despiking resulted in a significantly increased number of artefact-free trials (t=6.72, p<0.001) and fewer ICA ICs (t=9.6, p<0.001) needing to be removed (all paired t-tests). The number of rejected trials after despiking decreased from 8±3 (mean±std) to 3±2 (mean±std) and the number of removed ICs decreased from 24±6 (mean±std) to 12±4 (mean±std).

### Gamma responses to pain

Fig. 3 displays the effect of despiking on pain-related gamma activity. The maps show the typical topographical distribution with maxima at lateral and frontocentral electrodes. The comparison between both analysis strategies did not show any significant differences, neither for the comparison of induced gamma responses nor for the encoding of pain intensity (p<0.05, FDR uncorrected).

**Figure 2.**
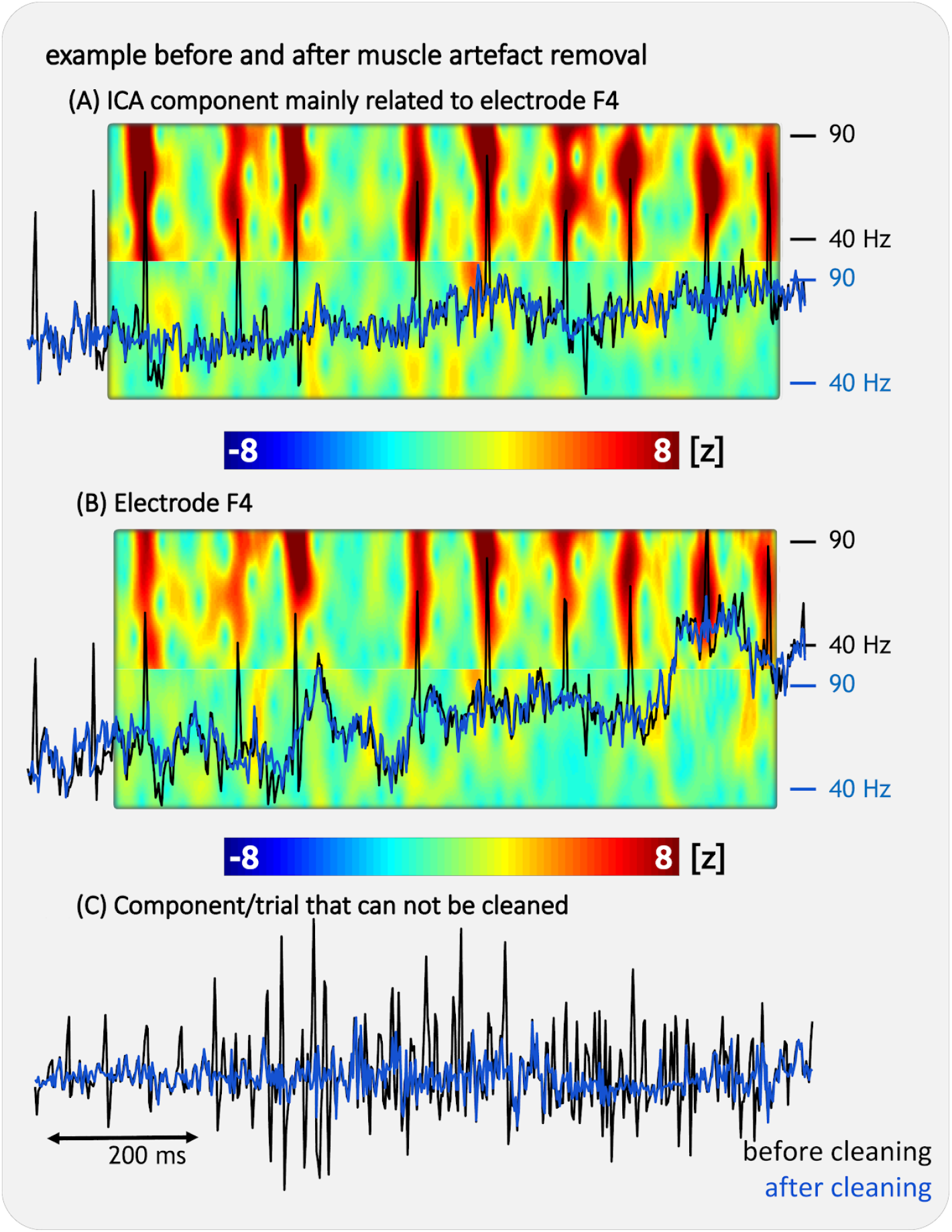
The figure shows an example of the result of the cleaning process. We applied the algorithm on ICA decomposed data. The component (A) has a topography with the highest weight from electrode F4 and mainly contains the high-frequency aspects of the data. The TFR of this selected IC time course shows the broad-band effect of muscle spikes before (2A, upper part) and after cleaning (2A, lower part). The lower part of the TFR shows the cleaned TFR window. (B) The TFR of this selected time course from electrode F4 shows the broad-band effect of muscle spikes before (2B, upper part) and after cleaning (2B, lower part). The signal at electrode F4 can result from the artefact removal of several ICs. The lower row (C) shows an example where the artefacts can be substantially reduced but the data can not be entirely cleaned. The trial or the entire IC need to be removed from the data.

**Figure 3.**
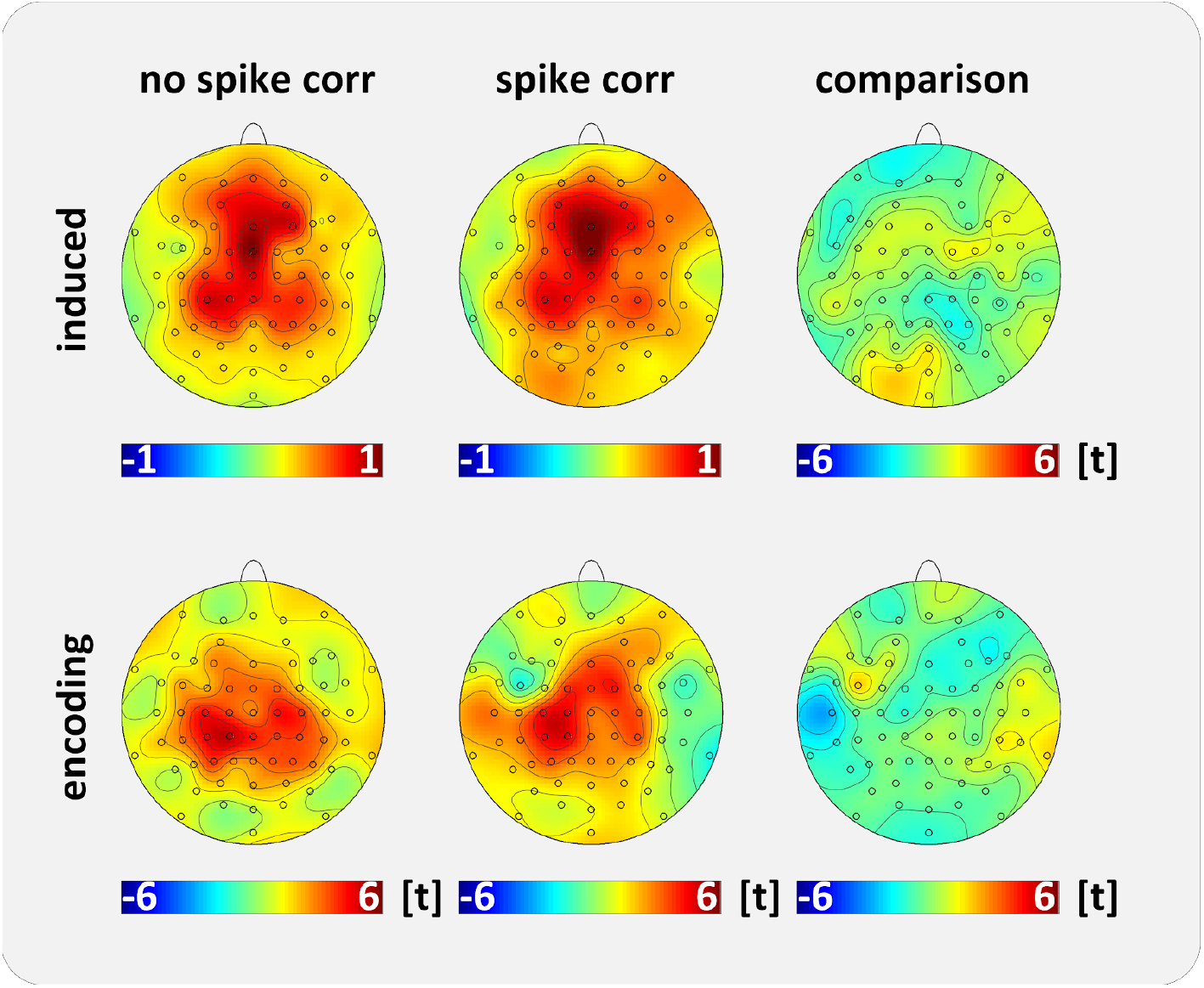
Topographical maps for pain induced gamma activity and the encoding of pain intensity by gamma responses. The comparison of both data cleaning approaches (with and without spike correction) did not show any differences.

## Discussion

The present study aimed to quantify the benefits of a novel algorithm to remove muscle spikes from EEG data. Three aspects need to be emphasised: *first*, the despiking tool allowed us to secure more data for the statistical analysis; we retained a greater amount of trials and ICs, thus supporting the successful performance of the tool. *Second,* the tool specifically removes muscle artefacts; the previously reported pain-related cortical effects in the gamma range were preserved after the application of the muscle despiking algorithm. *Third,* the validation of the despiking tool allows for the application to data where the removal of muscle artefacts is more challenging; this is the case for studies investigating gamma activity in resting-state EEG studies or in paradigms that investigate ongoing cortical processes, e.g. long-lasting (tonic or chronic) pain.

### Spike detection and removal

The despiking tool significantly improved the quality of our data and allowed us to preserve a high quantity of cortical information. We were relying on the work by Nottage and colleagues and utilised Gabor functions to fit and to remove single spikes (Nottage et al., 2013). Their algorithm specifically focuses on saccades and requires densely sampled data (>5kHz), without powerline noise contamination in the raw data. However, in order to remove not only saccades but all muscle spike artefacts, we followed a different path. Our approach does not require these prerequisites as it makes all muscle spikes (saccades, masseter, neck) more easily detectable by separating the underlying sources through ICA decomposition. We can distinguish three types of components:

> *First,* we observed nearly artefact-free components that would need minimal treatment. Here, we would only find occasional single spikes. A higher spatial sampling (>64 electrodes) could result in a better separation of the sources and would potentially yield completely artefact-free ICs that could be spared from artefact cleaning.
>
> *Second,* we identified highly artefact-contaminated ICs that are related almost entirely to muscle activity. With our 64 channel electrode array, single muscle spikes were often overlapping, which makes a sufficient fit with Gabor functions impossible. These ICs are mainly contributed by activity recorded from inferior temporal, fronto-temporal, and occipital electrodes (masseter or neck muscles sources). As a result, the underlying cortical signal is barely noticable. Hence, these ICs can not be sufficiently cleaned with the despiking tool and are suggested to be completely deleted from the data. A higher spatial sampling with an increased number of electrodes might help here; generating more independent ICs could potentially better separate overlapping muscle spikes. However, even without removing the component, the tool can scrub the largest spikes and therefore improve the SNR.
>
> *Third,* there is a continuous gradient between almost artefact-free ICs and highly contaminated ICs. Along the gradients there are ICs that exhibit occasional and distinct non-overlapping spikes. Such ICs would have been removed without a suitable treatment but can be spared from complete removal due to the application of the muscle artefact removal tool.

The ICA decomposition is particularly effective in cases with mixed cortical and muscular origin; the example in Figure 2 illustrates that not all muscle spikes recorded at electrode F4 originate from the same underlying source. The decomposition into components helps to single out the spike in the temporal domain and enables an optimal fitting with Gabor functions for relatively low sampling rates of 1000 Hz.

### Application to event-related EEG data

Most publications reporting the analysis of neuronal gamma activity used ICA to remove muscle activity from EEG signals (Hipp and Siegel, 2013). The fact that muscular electromagnetic signals originate from outside the electrically-resisting skull results in focal and high-magnitude muscle spikes, particularly at inferior EEG electrodes. Unlike cortical activity, there is no smearing of the cortical signal through skull transmission (Nunez and Srinivasan, 2006). These aspects make the different sources of muscular activity more separable and easier to detect. However, most studies rely on a limited number of electrodes, which in turn limits the number of separable and independent components.

Data analysts need to find a reasonable cut-off point to distinguish between “good” and “bad” ICs. This is relatively easy for event-related studies of pain (Chien et al., 2014) and other sensory modalities (Yuval-Greenberg et al., 2008). Evaluating IC topographies and averaged time-frequency decomposed trials in ICA space helps to distinguish between task-related neuronal activity and artefacts in most cases. Therefore, even a large number of rejected ICs would not compromise the estimation of cortical effects because the averaged TFR of the component shows whether there’s a stimulus-related cortical response. The current algorithm could be useful in uncertain cases, where no clear decision can be made about an IC as it may contain both cortical activity and muscle spikes.

### Application to continuous EEG data

The present analyses on event-related data show that the benefits of an additional spike removal, compared with a rigorous artefact removal without it, are barely noticeable. Our results suggest that a thorough data cleaning on a high number of electrodes (and a high number of separable sources) and a high number of (discardable) trials may not require additional spike removal, provided that the data analyst will follow (1) the recommendations of Muthukumaraswamy (Muthukumaraswamy, 2013), and (2) preserve the task-relevant processes by averaging epoched and time-frequency decomposed single trial ICA data, (3) as well as by removing artefact ICs and artefact contaminated trials.

In contrast, with resting-state studies on pain (Cao et al., 2016; Cauda et al., 2009; Dinh et al., 2019) or in studies investigating ongoing neuronal processes (May et al., 2019; Nickel et al., 2019; Schulz et al., 2015), we can not be entirely sure whether an artefact-contaminated IC also contains relevant cortical processes of interest. Without sufficient muscle artefact cleaning, findings can be falsely interpreted as originating from cortical sources. This can happen when ICs are spared from cleaning because they consist mostly of cortical activity; occasional muscle activity at task-relevant time frames may lead to false positives. Even IC time courses with a reasonable topography can contain occasional spikes whose shape suggests a muscular origin.

The present tool can increase confidence regarding the evaluation of ICs. Potentially important information can be secured which would otherwise have been lost. As an example, some head muscles (m. frontalis) are mainly projecting to fronto-central electrodes. Such artefact topographies with a fronto-central maximum are present in all participants. These artefact topographies, however, are perfectly overlapping with findings of the cortical encoding of long-lasting tonic and chronic pain (May et al., 2019; Schulz et al., 2015). The present algorithm can provide more confidence that findings on the encoding of long-lasting pain in the gamma range at frontocentral electrodes are exclusively caused by cortical processes.

### Limitations

There are a few limitations for the use of the tool that need to be considered. The entire process, and particularly the fitting of the Gabor functions, is very time consuming and can last a few hours on a high-performance multicore machine. Currently, the definition of required parameters needs to be done manually, as ICs contain various shapes of muscle spikes with different features, frequencies of occurrence, and spike densities. Future versions of the tool may define parameters through supervised machine learning approaches that are based on a large database. In addition, the tool may benefit from a faster programming language such as Julia, NumPy, or C++. The current application of the tool would have benefited from a high-density electrode array (>64 electrodes, with a higher density at artefact sites) to better separate the underlying sources of cortical and muscular activity. The tool does not make a careful inspection of single time-frequency decomposed trials redundant but limits the probability to preserve artefact-contaminated trials.

### Summary and outlook

By applying the tool to remove muscle spike activity, we obtain high data quality by preserving task-relevant gamma oscillations of cortical origin. We were able to precisely detect, gauge, and carve out single muscle spikes from the time course of neurophysiological measures. We speculate that this tool is particularly beneficial for studies with a less thorough cleaning procedure, whereby the majority of artefacts could have been eliminated.

We suggest the application of our validated tool for studies investigating gamma activity that contain a low number of trials as well as for data that are highly contaminated with muscle artefacts. We further recommend our tool for studies that investigate event-free continuous EEG recordings.

## Software availability

The muscle artefact tool is available on the website of the senior author (www.pain.sc).

## Conflict of interest

None of the authors declared any conflict of interest.

## Acknowledgements

We thank Dr Ulf Baumgärtner and Dr Rolf-Detlef Treede for their comments on the manuscript.

## Declarations of interest

none.

## Notes

### Competing Interest Statement

The authors have declared no competing interest.

